# Directed cell migration is a versatile mechanism for rapid developmental pattern formation

**DOI:** 10.1101/2025.07.24.666657

**Authors:** Chengyou Yu, Malte Mederacke, Roman Vetter, Dagmar Iber

## Abstract

The evolution of multicellular organisms hinges on self-organization mechanisms that generate tissues with diverse functions. A central process is the breaking of symmetry to form spatial patterns from initially uniform conditions. Among the various mechanisms proposed, directed cell migration—driven by chemotaxis, durotaxis, differential adhesion or other processes—offers a compelling strategy to organize tissues rapidly and robustly. Here, we unify these concepts into a general mathematical framework and show that it can produce diverse spatial patterns across one-, two-, and three-dimensional domains. Using numerical simulations and stability theory, we characterize the emergence, geometry, and formation speed of these patterns. Our findings provide a mechanistic understanding of morphogenesis beyond the traditional chemical or mechanical patterning paradigms and offer a quantitative foundation to guide pattern formation in tissue engineering and regenerative medicine.

## Introduction

Multicellular development is orchestrated through a series of self-organizing processes that generate robust spatial patterns from initially homogeneous conditions. Classical explanation attempts for pattern formation can broadly be categorized into three families of concepts: chemical reaction–diffusion systems (most famously, the Turing mechanism [1], the French Flag model [2] and traveling waves [3]), mechanical instabilities driven by differential growth and tissue buckling [4–8], and cellular motion such as sorting and migration [9–11].

Reaction–diffusion systems provide a powerful framework for explaining morphogenetic patterning, but some act too slowly to account for rapid pattern emergence across large domains [12], and identifying the Turing pair has not always been an easy task [13–15]. In addition to the requirement of at least one—if not two—free chemical reactant interacting with the tissue, Turing’s patterning mechanism works only within a kinetic regime (the Turing space) that is typically narrow [16] and the resulting patterns can have a high qualitative sensitivity to initial noise levels [17].

Mechanical buckling due to differential growth can quickly yield periodic structures [18], but requires room for the tissue to deform out of plane, which is not always a desired patterning outcome. Unless localized by some form of pre-patterning or spatial constraint, large-scale tissue deformations ensue. While not depending on a chemical signal per se, mechanical buckling requires some form of spatial inhomogeneity to begin with, such as at least two layers of tissue with different mechanical properties and growth patterns to produce undulations [3, 19]. This mechanism is thus not easily applicable in more uniform tissue environments such as a simple epithelial sheet.

But Nature has found a way to pattern tissues without additional chemical signals or changing their shape, using their very own building blocks—the cells—as agents: Through a combination of random and active directed motion, cells can agglomerate and sort [20, 21] within the tissue itself. In recent years, more and more examples of fast developmental processes that involve considerable cellular movement have been identified, including tracheal cartilage ring formation [22], hair follicle patterning [23, 24], and the rearrangement of patchy domains of Wnt activity into a single pole that defines the anterior–posterior axis in gastruloids [25–27]. Cells can interpret a range of environmental signals—chemical gradients (chemotaxis) [28, 29], adhesion landscapes (haptotaxis) [30], substrate mechanics (durotaxis) [31], and physical topography (topotaxis) [32]—to guide their migration.

One key driver is differential cell adhesion (DCA) [9, 33], which has been studied through both discrete cell-based approaches and continuum models. Discrete models capture individual cell behavior in fine detail [34], but they often struggle to scale to biologically relevant tissue sizes. In contrast, continuum models allow for efficient simulation of large-scale tissue dynamics and provide analytical tractability [35–38], making them particularly useful for studying macroscopic pattern formation. Mathematically, these continuum models are expressed in the form of partial integro-differential equations (PIDEs). Their use in cell biology and dates back to the late 1990s [39] but is predated still in ecology (see [40] for a brief review). PIDEs are typically challenging to solve due to their hybrid form and require tailored numerical discretizations and algorithms [41–43]. Previous DCA studies based on PIDEs employed custom finite-volume solvers and left questions regarding dimensionality, scope, and parameter dependency open. The requirement for expert knowledge to simulate patterning with continuum DCA models may partially account for the under-appreciation of their potential in explaining developmental pattern formation.

Here, we study a unified model of directed cell migration (DCM) that generalizes classical DCA. We implement it in a common general-purpose finite element solver, which makes it accessible at scale. Combining this framework with linear stability theory, we explore the conditions under which spontaneous symmetry breaking leads to pattern formation in tissues across one, two, and three dimensions, both in static and growing domains. Our results show that patterns driven by DCM can emerge within short, biologically relevant timescales, and we identify conditions under which these patterns are robust to noise and geometric constraints. Furthermore, we demonstrate how patterns can be oriented through anisotropic behavior, offering a strategy for controlled directional patterning. By establishing a general and extensible framework for directed cell migration, our work bridges the gap between individual cell behavior and tissue-scale morphogenesis. As a versatile mechanism for rapid tissue patterning, DCM has the potential to join the established chemical and mechanical models as a major class of models for developmental pattern formation. Our findings thus offer a new perspective on developmental processes and provide a quantitative tool that may guide pattern formation in regenerative medicine, tissue engineering, and organoid design.

## Results

### Continuum Model of Directed Cell Migration

Unifying previous continuum descriptions of DCA [36–39, 44], we model pattern formation through the migration of a cell population within a tissue or matrix environment described as a convection-diffusion-like system of the general form

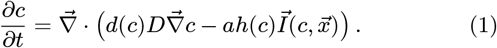

Here, 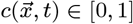 represents the normalized concentration of cells (or similar agents, but we refer to them as cells here) at position 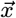 within the domain of interest at time *t* (Fig. 1A). Depending on the context, this may be the concentration of a single cell type within a solution or matrix environment relative to a certain capacitance, or the partial concentration of a certain cell type within a heterotypic tissue. *c* may also represent the fraction of (un)differentiated cells in a ensemble of stem cells. The remaining proportion 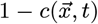 then represents the sum of media and other cell populations.

**Figure 1:**
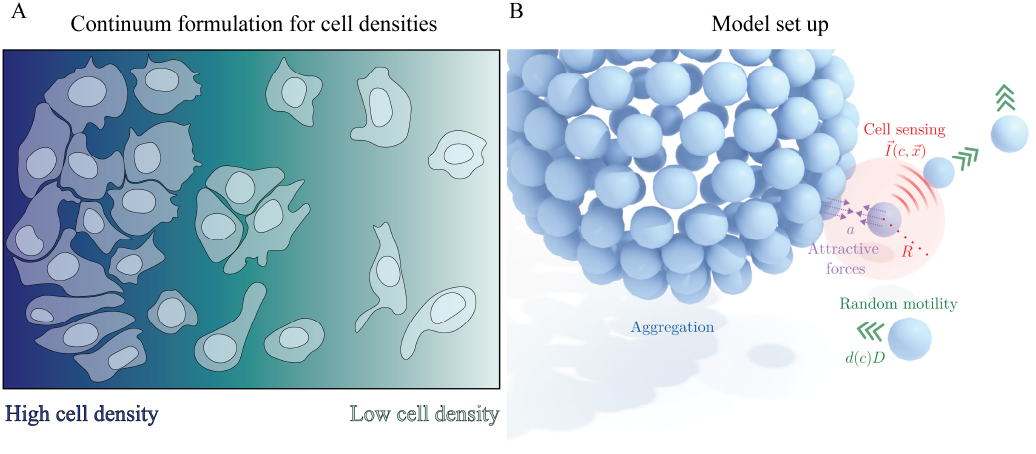
Schematic illustration of the continuum cell density models and the directed cell migration model. **(A)** Continuum formulations describe local cell density, instead of representing individual cells as agents. An area with more cells and/or a tighter packing carries a higher cell density (darker blue color). **(B)** In the DCM, cells aggregate due to attractive directed forces between them within a cell sensing of spatial range *R* (red circle). Cells migrate actively in response to these forces, but may also move due to random motility or passive dispersal.

The transport model is divided into two contributions in the divergence operator, a passive non-directional dispersive term and an active directed migratory term. The first term, 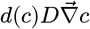, describes random cell motility or diffusion with coefficient *d*(*c*)*D*. In its simplest form, *d*(*c*) = 1 such that the diffusivity is a constant. *d*(*c*) = *c* corresponds to advection by a populationpressure-driven velocity field [37], which lets cells move faster the larger the local density. Cells in the developing chicken limb mesenchyme, for instance, indeed move slightly faster in pre-cartilage condensations than in non-condensed regions [45]. Feedback mechanisms may lead to other functional relationships for *d*(*c*), reflecting other physical or biological bases for cellular dispersal.

The second term in the divergence operator models directional locomotion with activity strength *a*, a modulator *h*(*c*), and a direction functional 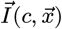, which is not necessarily local. It may be interpreted as a convective term, especially if *h*(*c*) = *c*, for which it becomes 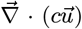 with velocity field 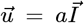. Various ways to define the migratory direction 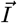 that lead to patterning have been proposed in the literature, some based on local information of stress gradients [24, 46], others through the notion of chemical or mechanical communication within a certain local neighborhood of a cell [36–38]. Here we use the latter, somewhat more general and potentially more directly biologically interpretable integral form, turning the model into a PIDE:

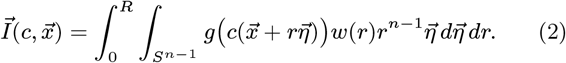

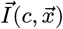 integrates some functional readout of the local concentration, *g*(*c*), multiplied by a unit directional vector 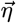 and a dimensionless weighing function *w*(*r*) that describes the dependency of the migratory cue on the distance between cells, within an spherical neighborhood of radius *R* in an *n*-dimensional domain. *S*^*n−*1^ denotes the (*n−* 1)-sphere. Note that in this form, the directional information is isotropic, since the integral does not include a directional bias. Anisotropic extensions that weigh different directions differently, or integrate over a nonspherical neighborhood, are also possible in principle, but are not included in our analysis here.

Note that this description is not limited to adhesive forces between cells, but covers attractive forces within a range of *R* in general. The model thus features random motility, cell sensing and attraction (for example through cytonemes or filopodia of maximum length *R*) and thus, aggregate formation (Fig. 1B). Different model variants can be built by choosing specific forms for the modulator functions *d*(*c*), *g*(*c*) and *h*(*c*). *g*(*c*) and *h*(*c*) may typically either be *c* or *c*(1− *c*), where the latter theoretically limits cell concentrations to be bounded by a carrying capacity.

We non-dimensionalize Eqs. 1 and 2 by setting 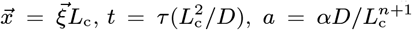, and *R* = *ρL*_c_, where *L*_c_ is a characteristic length scale. In most cases, when the sensing radius *R* is not varied, we conveniently choose *L*_c_ = *R*, i.e., *ρ* = The concentration *c* here is already normalized (*c* ∈ [0, 1]) as it represents a populational fraction. The non-dimensionalized equation reads

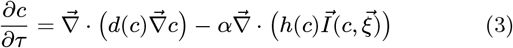

where

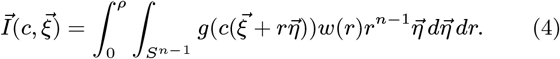

The dimensionless computational domain then has an edge length that we denote by *L*_0_. Unless specified otherwise, *L*_0_ = 10 is used in the following.

### Modulating the Patterning Phenotype

To quantify how the patterning phenotype dynamically depends on chosen model parameters, we implemented the DCM numerically with a commercial finite element solver (Methods) and performed simulations of a sample model assuming *d*(*c*) = 1, *h*(*c*) = *c*(1*− c*) and *g*(*c*) = *c* with different parameter settings in 2D and 3D under periodic boundary conditions, to imitate the patterning in large tissues (Fig. 2A). The reasons for the choice of this model will be explained later. The results show that the initial concentration has a considerable impact on the pattern (Fig. 2B). Patterns generated from low initial concentrations (e.g., *c*_0_ = 0.3) appear as individual and relatively spherical spots, and are almost inversions of patterns generated from high initial concentrations (e.g., *c*_0_ = 0.7) in this model variant. For *c*_0_ = 0.5, transient labyrinthine patterns emerge in an earlier developmental phase (*τ* = 10). Patterns tend to exhibit sharper boundaries with higher attraction strength values *α*. Similar patterning behavior could be found in 3D simulations as well (Fig. 2C).

**Figure 2:**
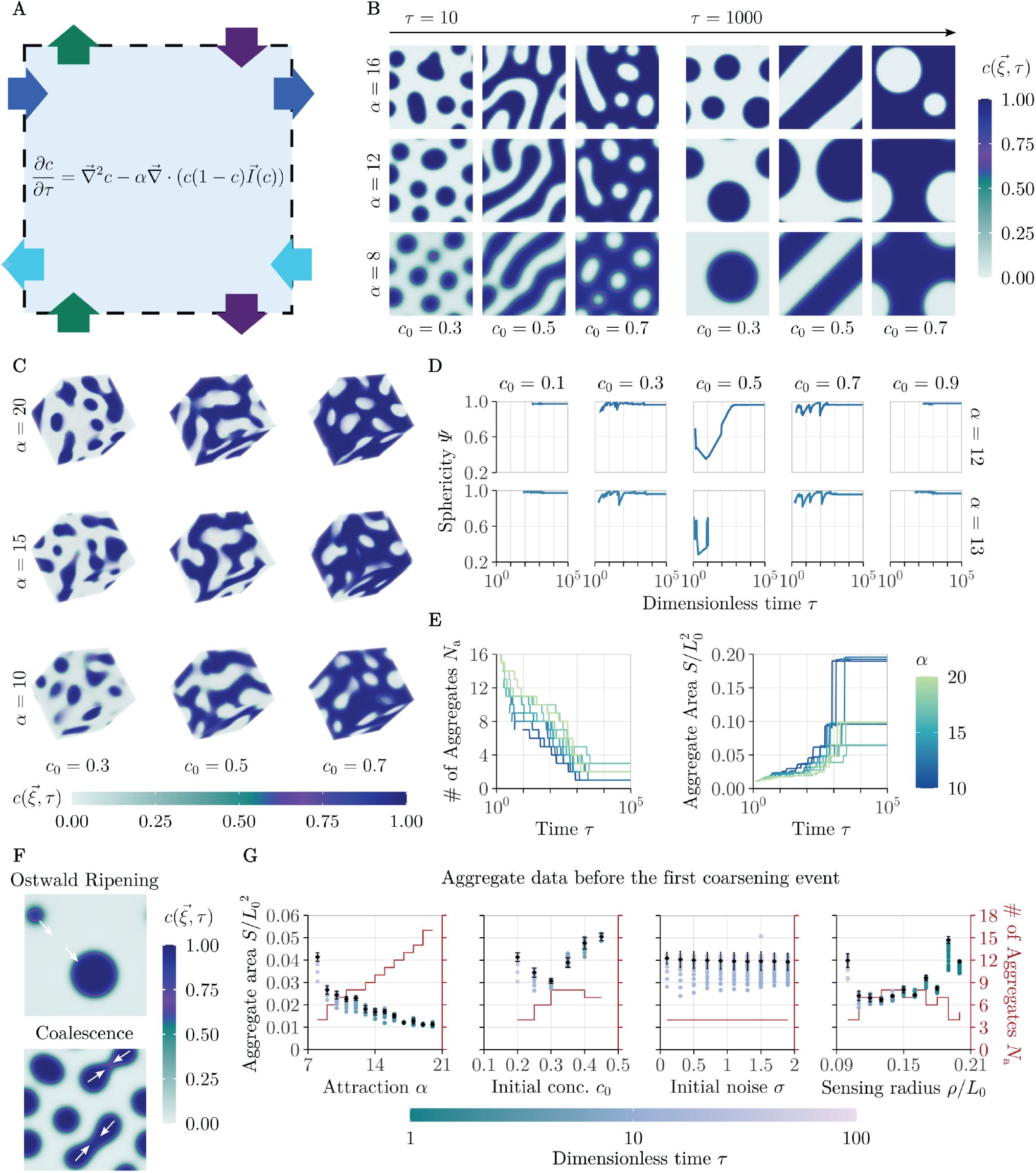
Pattern morphology under periodic boundary conditions. **(A)** Schematic illustration of the simulation settings. Simulations are run with periodic boundary conditions in the model variant with *d*(*c*) = 1, *h*(*c*) = *c*(1 *− c*), and *g*(*c*) = *c*. The blue shaded region represents the computational domain. **(B)** Patterns forming under different attraction strengths *α* and initial concentrations *c*_0_ at dimensionless times *τ* = 10 (left) and *τ* = 1000 (right) in a 2D domain. **(C)** Patterns forming under different *α* and *c*_0_ at *τ* = 20 in a 3D domain. **(D)** Evolution of the average sphericity Ψ of aggregates over time under different *α* and *c*_0_. **(E)** Changes of the number of aggregates *N*_a_ (left panel) and the average relative area of aggregates *S/L*^2^ (right panel) over time for different *α* values. *L*_0_ = 10 is the dimensionless side length of the simulation domain. Initial concentration: *c*_0_ = 0.2. **(F)** Examples of coarsening by Ostwald ripening (top panel), where an aggregate shrinks and eventually dissolves into other aggregates, and coalescence (bottom panel), where multiple nearby aggregates merge into a single cluster. **(G)** Average sizes and numbers of aggregates between the time point of patterning and the first Ostwald ripening or coalescence event for (from left to right): 1. different *α* values (*c*_0_ = 0.2), 2. different *c*_0_ values (*α* = 8), 3. different *σ* values (*α* = 8, *c*_0_ = 0.2), and 4. different *ρ* values (*α* = 8, *c*_0_ = 0.2). Colored points indicate the average size of aggregates at a single time point. Black diamonds and error bars indicate the mean *±* SD across all time points. The brown lines represent numbers of aggregates.

As time progresses, aggregates forming at a lower *c*_0_ typically show coarsening by two behaviors: They may shrink gradually and eventually dissolve into other, larger aggregates, a process known as Ostwald ripening [47] (Fig. 2F, top panel), or they may coalesce with nearby aggregates to form larger clusters (Fig. 2F, bottom panel). Inverse aggregates (gaps) formed at *c*_0_ *>* 0.5 show analogous behavior. This is further demonstrated in Fig. 2E, where we tracked the aggregate number and average aggregate size over time for *c*_0_ = 0.2 under different *α* values. As expected, aggregates decreased in number but grew over time. Furthermore, systems with higher attraction strength tend to produce more but smaller aggregates.

The sphericity 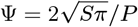 (in 2D), where *S* and *P* are the area and perimeter of the aggregates, is mainly dependent on the initial concentration of the system (Fig. 2D). We observed higher average sphericity early on in systems with initial concentrations further away from *c*_0_ = 0.5. Except for *c*_0_ = 0.5, the sphericity tends to increase over time and asymptotically approaches unity, i.e., the aggregates form perfect spheres. Intermediate drops in sphericity indicate coalescence events, as coalescence transiently produce deformed spheres. For the symmetric initial level *c*_0_ = 0.5, a variety of patterns is possible after long enough time. A selection is shown in Fig. 2B at *τ* = 1000, such as diagonal stripes (*α* = 8, 16), and spherical spots (*α* = 12). Therefore, in cases such as *α* = 12, the sphericity approaches unity over time, whereas in cases like *α* = 13 (yielding a horizontally domain-spanning stripe, not shown here), the sphericity cannot be defined after a certain time point (here at *τ ≈* 10) as the aggregates extend infinitely (Fig. 2D).

To investigate the impact of parameters on the size and number of the condensates, we measured them within the time frame starting from the time point of pattern formation *τ*_p_, until the first coarsening event to eliminate the effect of these events as confounding factors (Fig. 2G). Aggregate sizes monotonically decrease with increasing attraction strength *α*, while the number of aggregates steadily increases, in agreement with Fig. 2E. In contrast, the initial concentration *c*_0_ has a non-trivial quantitative effect on the pattern, with a local minimum in the aggregate size and a local maximum in their number at *c*_0_ ≈ 0.3. The initial random noise level *σ* (Methods) does not have a measurable effect on the aggregate size. As the sensing radius *ρ* is varied, the average size of aggregates first drops from *ρ/L*_0_ = 0.1 to *ρ/L*_0_ ≈ 0.13, but then rises again as *ρ* is increased to *ρ /L*_0_ = 0.2. For all four parameters, aggregate size and number are inversely related, such that a decrease in size is accompanied by an increase in quantity.

### Pattern Formation

#### Critical Conditions

Patterns emerge only under certain conditions that can be mathematically determined through stability analysis. We performed a linear stability analysis similar to that of Jewell et al. [48] to find the conditions for pattern formation based on the system parameters and model variants (Methods). In 2D, assuming *g*′(*c*_0_) *>* 0, the conditions for pattern emergence from a uniform but noisy initial concentration *c*_0_ can be summarized as

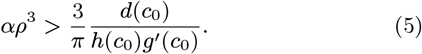

Eq. 5 shows that the attraction strength *α* and the sensing radius *ρ* will have to exceed certain thresholds for patterns to form, and they can mutually compensate for one another. Similar conditions apply in 1D and 3D, with an exponent *n* + 1 for the dimensionless sensing radius *ρ* in *n* dimensions (Methods). Thus, weaker attraction (smaller *α*) needs to be compensated by a larger sensing region according to *ρ ~ α*^*−*1*/*(*n*+1)^ to obtain patterning, which becomes ever easier in higher dimensions. The effect of the initial concentration *c*_0_ on patterning depends on the modules *d*(*c*_0_), *g*(*c*_0_), *h*(*c*_0_), and can both shrink or widen the patterning space.

To better visualize the constraints on *c*_0_ in different model variants, we chose three representative models: Model 1: *d*(*c*) = 1, *h*(*c*) = *c, g*(*c*) = *c*(1 *− c*), Model 2: *d*(*c*) = 1, *h*(*c*) = *c*(1 − *c*), *g*(*c*) = *c*, and Model 3: *d*(*c*) = *c, h*(*c*) = *c*(1 − *c*), *g*(*c*) = *c* (Fig. 3A). For a fixed sensing radius *ρ* = 1, *α* must exceed a minimal threshold independent of *c*_0_ as a necessary (but generally not sufficient) condition for patterning, which can be found by minimizing *d*(*c*_0_)*/*(*h*(*c*_0_)*g*′(*c*_0_)). Depending on the functional form of the three modules, this minimum is attained in the interior of the initial concentration range *c*_0_ ∈ [0, 1] or at a boundary.

**Figure 3:**
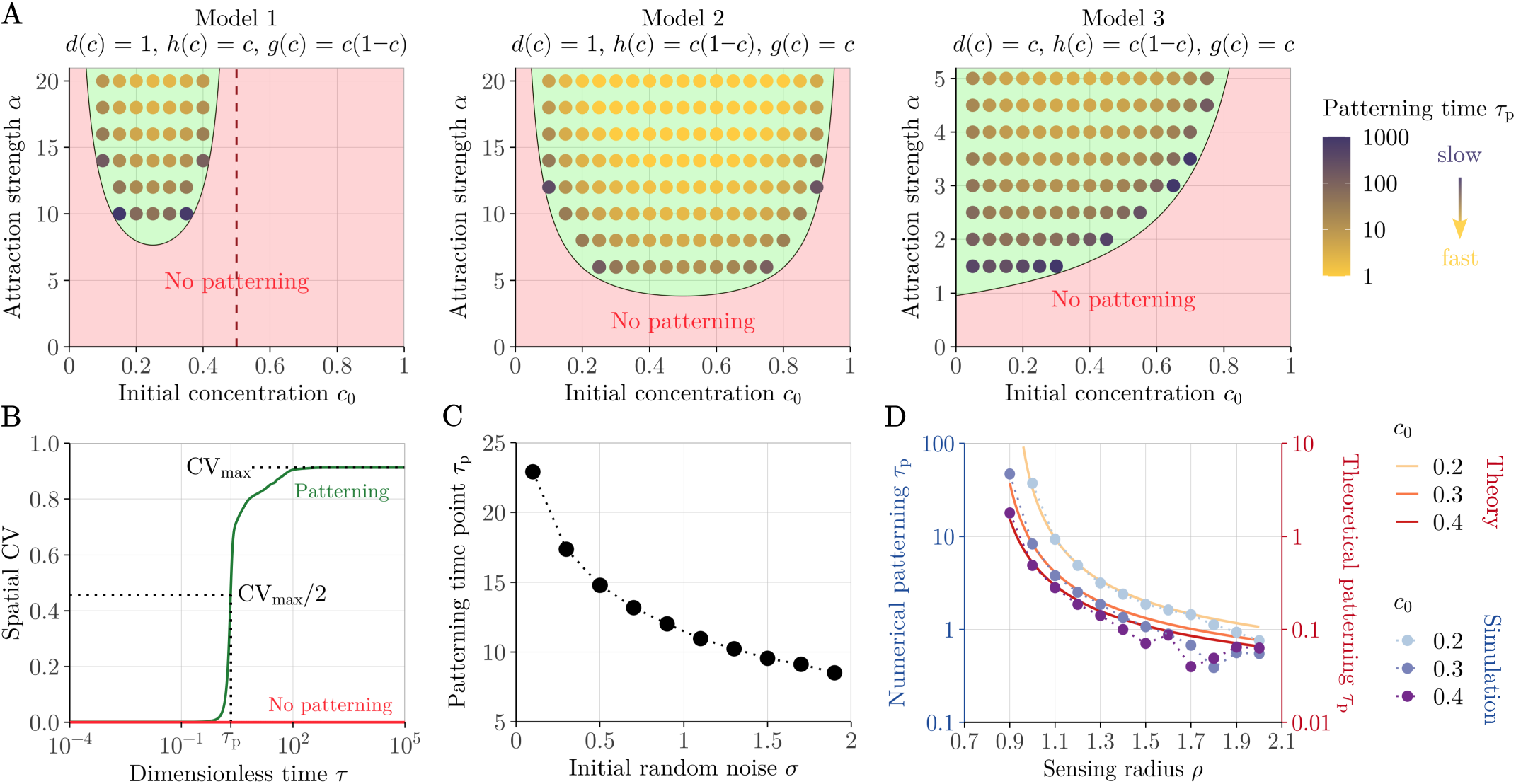
Patterning conditions and speed. **(A)** Phase space of patterning for three representative model variants. Green-shaded areas indicate *α >* 3*d*(*c*_0_)*/*(*h*(*c*_0_)*g*′(*c*_0_)*πρ*^3^). Points represent numerical simulations that produced patterns; absence of a point in the grid indicates no pattern. Colors of points represent the time point of pattern formation, *τ*_p_, as defined in (B), from slow (dark purple) to fast (yellow). Simulation settings: *ρ* = 1, *σ* = 0.01. **(B)** Definition of the patterning time *τ*_p_ as the time point at which 50% of the maximum spatial coefficient of variation (CV) of concentration across the computational domain at *τ* = 10^5^ is reached. The green and red curves show representative example cases with and without pattern emergence. **(C)** Influence of the initial random noise level *σ* (Methods) on the patterning time. Simulation settings: *ρ* = 1, *α* = 8, *c*_0_ = 0.2. **(D)** Influence of the sensing radius *ρ* on the numerical (blue color scale) and theoretical (red color scale) patterning time *τ*_p_ for different *c*_0_ values. Simulation settings: *α* = 8.

For random diffusion-based models (*d*(*c*) = 1, Models 1 & 2), the minimum is located at the value of *c*_0_ for which *g*′(*c*_0_)*h*′(*c*_0_) = − *g*^*′′*^(*c*_0_)*h*(*c*_0_). For Model 1, this is at *c*_0_ = 0.25, for Model 2 at *c*_0_ = 0.5. Those are the initial concentration levels for which patterning requires the least minimal attraction strength and smallest sensing radius, i.e., for which it may be easiest to achieve biologically. Beyond that minimum threshold, the patterning condition depends on the initial concentration *c*_0_. The higher the attraction strength between cells, the wider the range of initial concentrations that allows for patterning.

Interestingly, the patterning condition (Eq. 5) has a pole at *c*_0_ = 0.5 for Model 1. Linear stability analysis (Methods) suggests that while patterning with *g*(*c*) = *c*(1*− c*) at *c*_0_ *>* 0.5 is possible in principle in 2D and 3D but not in 1D, it would require very large (and possibly biologically unphysiological) attraction strengths. We indeed find this confirmed in numerical simulations in 2D, where for the entire range of numerically feasible values attraction strengths (*α <* 20 at *ρ* = 1), patterns were observed only if the initial density of active cells was below one half (Fig. 3A, left). A fourth model variant with capacitance limitation in both *g*(*c*) and *h*(*c*) behaves analogously (Supplementary Fig. S1, left).

For a model with pressure-driven motility (*d*(*c*) = *c*, Model 3), the picture is qualitatively different. The critical patterning boundary monotonously increases in *c*_0_, such that the minimum lies at *c*_0_ = 0 (Fig. 3A, right). The biophysical requirements for patterns to form are thus lowest when only few actively migrating cells are present. We also tested further model variants and found analogous behavior (Supplementary Fig. S1, middle & right).

To further deepen our analysis, we selected Model 2, as it arguably offers the most interesting patterning space across a wide range of initial densities, but unlike the pressure-driven variants (*d*(*c*) = *c*), it does not produce patterns in the dilute active cell limit (*c*_0_ → 0). That a minimum threshold of contractile cells is required for phase separation has been noted also for related models [49]. We thus use *d*(*c*) = 1, which, additionally, is numerically more benign.

#### Patterning Speed

To determine the speed of patterning, we defined the patterning onset *τ*_p_ as the time point at which the spatial coefficient of variation (CV) of concentration across the computational domain reaches 50% of the maximum steady-state value CV_max_, which is considered attained at *τ* = 10^5^ (Fig. 3B, dark green curve). This method is robust as the rise of the CV around *τ*_p_ is steep compared to other time points. In the case of no patterning, the CV stays at a low level comparable to the initial value (red curve).

The timing of pattern formation is mainly modulated by four factors: The attraction strength *α*, the sensing radius *ρ*, the initial random noise level *σ*, and the initial concentration *c*_0_. By considering the inverse of the perturbation growth rate *ω* of the fastest-growing spontaneous patterning mode *k*_max_ in a linear stability analysis (Methods) [24], one can acquire a theoretical estimate of *τ*_p_:

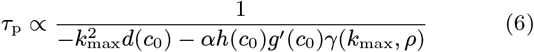

where *γ*(*k, ρ*) is a dimensionality-dependent function (Methods). The theoretical patterning time point is indeed proportional to the numerically observed one with a proportionality factor of about 10 (Fig. 3D, Supplementary Fig. S2). However, the numerical patterning time deviates moderately from theoretical expression (Eq. 6) in the strong activity regime: At large *α* and *ρ* values, and for *c*_0_ close to 0.5 in Model 2, numerically stable simulations are challenging.

Pattern emergence generally accelerates the further into the patterning space the system parameters lie (Fig. 3A). With greater attraction strength *α* comes faster cell migration. Increasing the initial noise level *σ* also accelerates pattern emergence (Fig. 3C). Noisier initial distributions will lead to locally larger 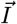 values, resulting in faster cellular migration. Varying *σ* across its entire range (0, 2) only enhances the speed about 2.7-fold, suggesting that the patterning time point is less sensitive to initial noise than to other parameters such as *c*_0_, for which an increase from 0.2 to 0.5 will result in about an 8.5-fold boost in patterning speed under the same *α* and *ρ*. Larger sensing radii also accelerate the pattern formation (Fig. 3D), since cells with a greater sensing range can detect crowded regions more swiftly and direct their movement toward them, instead of relying on diffusion to bring them in their vicinity. The final factor impacting the pattern formation time is the initial concentration *c*_0_, which has a non-monotonous effect, as the existence of a limiting capacity will set a maximum patterning speed, as indicated by Eq. 6. Raising *c*_0_ from zero will first lead to quicker patterning, but will then delay the patterning onset once *c*_0_ exceeds a certain level (Fig. 3A) due to overcrowding.

#### Patterning in Finite Tissues

The aforementioned numerical analyses were performed under periodic boundary conditions (PBC). To explore patterning in finite tissues, we carried out similar simulations under zero-flux boundary conditions in a bounded domain, assuming a vanishing concentration outside of the domain, which restricts intercellular attraction to within the tissue considered. This also guarantees that the *n*D spherical integral is well-defined at domain edges (Fig. 4A). We observed considerable boundary effects on the formation of aggregates, be it at low or high initial concentration levels (Fig. 4B). High-density aggregates generally tend to avoid the tissue boundaries and align in parallel to them. At *c*_0_ = 0.5, concentric ring-shaped structures formed, whereas lower or larger *c*_0_ produced spotted patterns away from the boundaries. Like in the case with PBC, pattern inversion was observed with respect to a reflection of *c*_0_ about 0.5, although the dependency is not perfectly symmetric. Over time, aggregates also coarsened (Fig. 4B).

**Figure 4:**
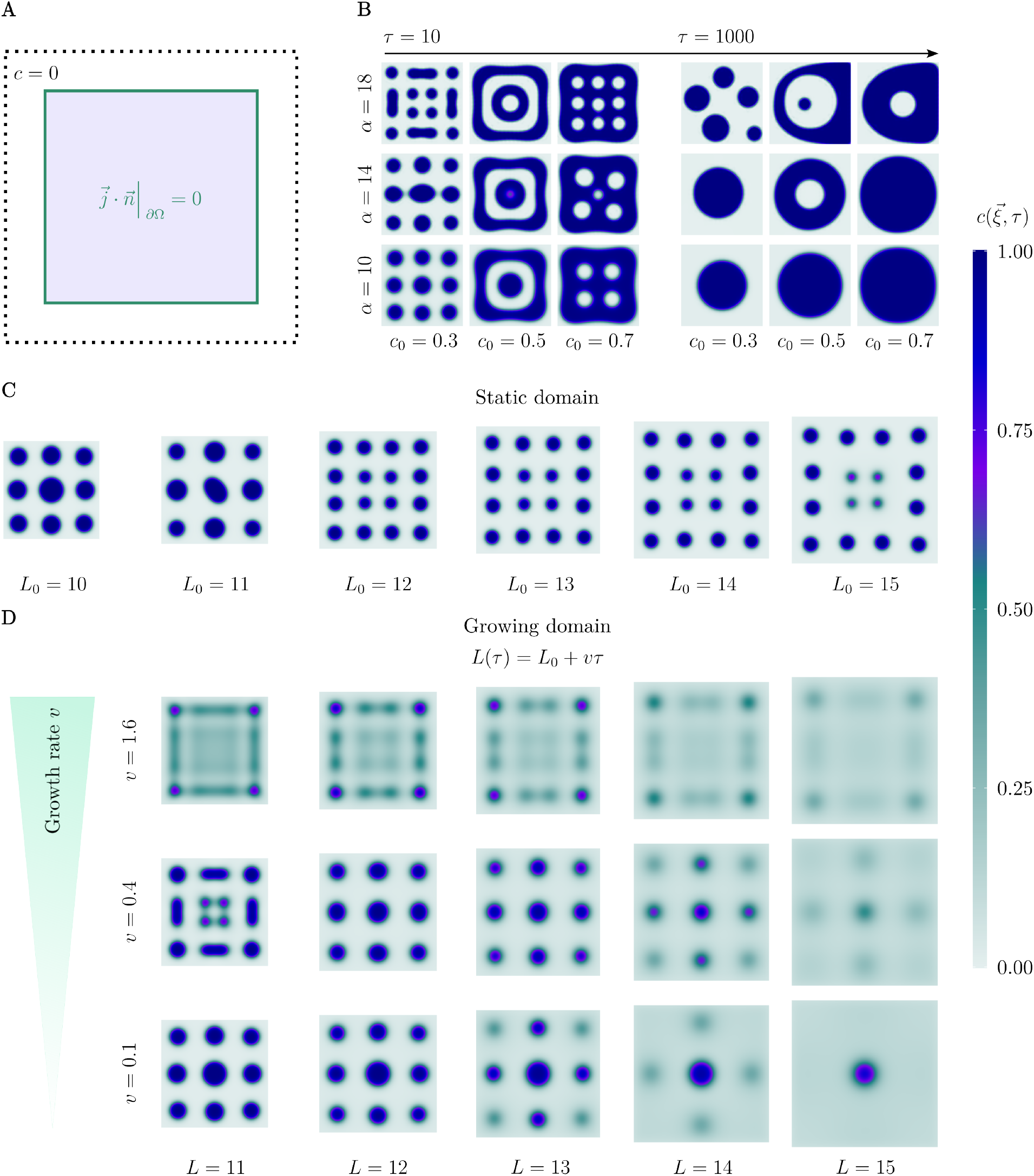
Patterning under zero-flux boundary conditions in a 2D domain. **(A)** Illustration of the simulation settings. The purple-shaded box is the computational region Ω. The dark green frame indicates zero-flux boundary conditions, mathematically described by 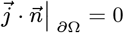, where 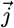 is the flux and 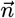 the local unit normal vector of the boundary *∂*Ω. The region outlined with dotted lines outside of Ω is not included in the computational domain, but has defined concentration *c* = 0 to assure that the spherical integral 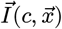 is well-defined everywhere. **(B)** Patterns formed under different *α* and *c*_0_ values at *τ* = 10 (left) and *τ* = 1000 (right). **(C)** Simulations with identical initial active cell mass *M* = ∫ _Ω_ *c*_0_*d*Ω = 30 spread within 2D square domains of different side lengths *L*_0_ at *τ* = 10, *α* = 12. **(D)** Representation of a growing domain (side length *L*(*τ*) = *L*_0_ + *vτ* changing from *L*_0_ = 10 to 15) under various growth rates *v* at different stages, for *α* = 12.

The size of the tissue influences the pattern morphology. We ran simulations in six static square domains with side length ranging from *L*_0_ = 10 to 15, containing the same active cell mass *M* = ∫ _Ω_ *c*_0_*d*Ω = 30. In our DCM model, dilution can lead to morphological thinning and spatial separation in parallel. We found more aggregates forming in larger domains, separated by larger gaps (Fig. 4C).

During development, tissues can grow in size. To capture the dynamic effect of dilution during domain expansion, we also modeled an isotropically growing square domain whose side length linearly increases over time at a growth rate *v*. To avoid computationally expensive remeshing, we mapped the problem from the Eulerian to a Lagrangian description (Methods), which allows the growing tissue to be simulated on a static domain.

We ran simulations starting from an initial side length of *L*_0_ = 10 growing up to 15 at varying growth rates *v* (Fig. 4D), and compared them to simulations on respective static domains (Fig. 4C). The cell aggregates tended to shift in position, aligning with the directions of domain expansion. They diluted with passing time, and tended to dissolve into neighboring aggregates through Ostwald ripening. With higher growth rates (e.g., *v* = 1.6), the time for nearby aggregates to coalesce before being separated by dilution decreased, resulting in early transient patterns which resembled those that developed in a static domain with longer side length (e.g., *L*_0_ = 15, Fig. 4C). Vice versa, for slower growth (e.g., *v* = 0.1), patterns resembled those developed in a smaller static domain (e.g., *L*_0_ = 10, Fig. 4C). These results demonstrate that the emerging patterns are strongly affected by early dynamics.

#### Anisotropy impacts pattern morphology

The patterns shown so far are mainly spherical and not oriented in any specific direction unless transient or impacted by boundary effects. In biology however, oriented patterns are common, and some have important functions for the viability of the organism, such as C-shaped cartilage rings surrounding and supporting the trachea [22] or the stripes in the fur of zebras. For these oriented patterns to emerge in the DCM model, some sort of anisotropy must be introduced. Here, we discuss two different approaches to orientate patterns: 1. anisotropy in the active cellular migration (attraction) itself, and 2. an anisotropically expanding attraction zone within the tissue.

##### Attraction anisotropy

The first type of anisotropy we consider is bias in the migratory strength in one of the spatial directions by a factor of *β ≥* 1 (Fig. 5A). This type of direct manipulation of model parameters to introduce orthotropy has previously been used to study directed stripe formation with Turing patterning, for instance [50, 51]. Mathematically, the anisotropic model can be written as

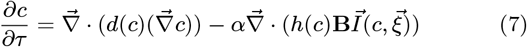

where **B** = diag(1, …, *β*) is a *n ×n* diagonal matrix and *β* ≥ 1 could be placed in any entry on the diagonal depending on the direction in which the patterns should be aligned.

**Figure 5:**
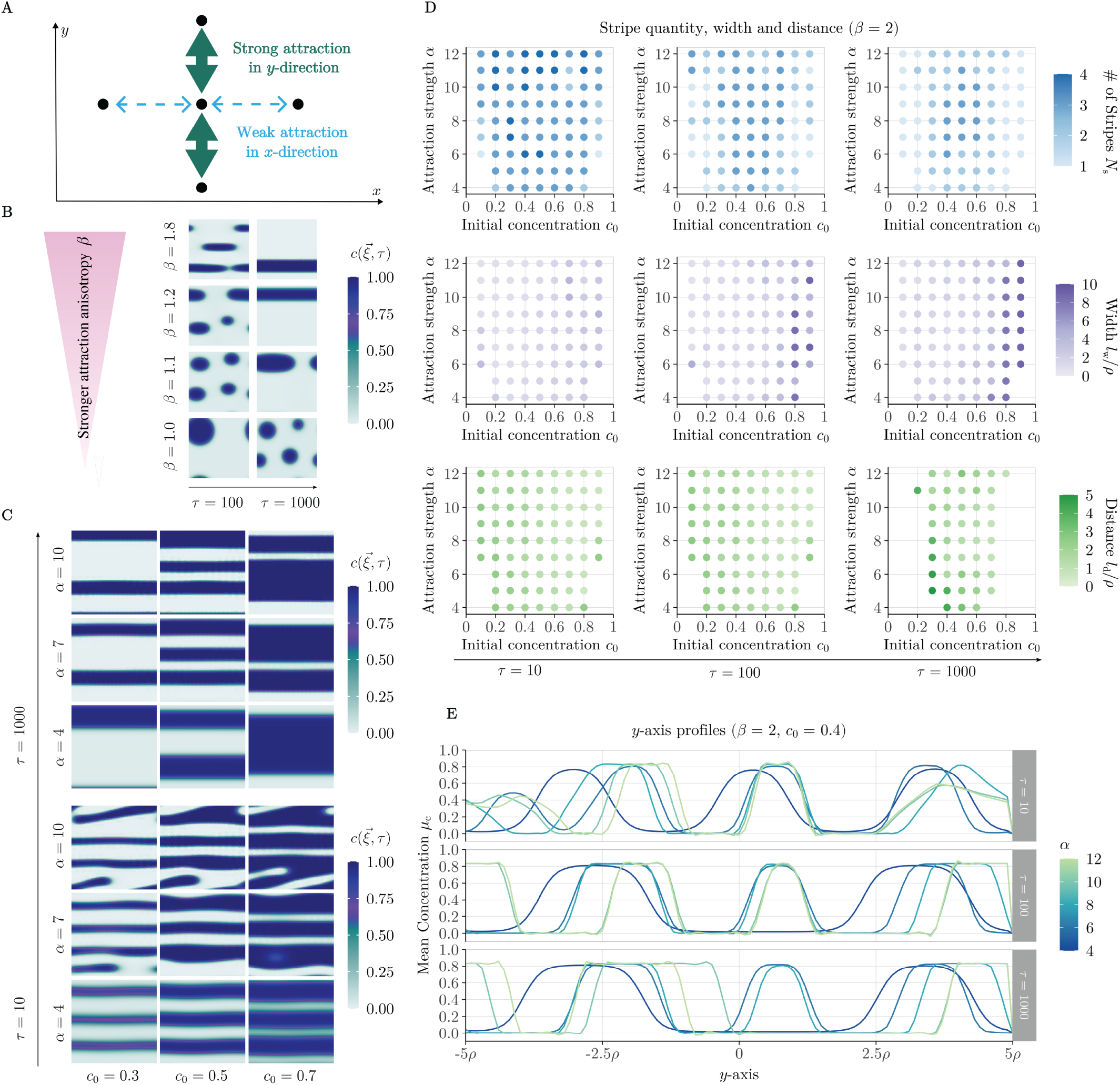
Anisotropic attraction strengths result in oriented patterns. **(A)** Schematic illustration of anisotropy in attraction strengths in different directions. We implement strong attraction in the *y*-direction and weak attraction in the *x*-direction. **(B)** Impact of the attraction bias *β* on patterning morphology at time points *τ* = 100 (left) and *τ* = 1000 (right). Simulation parameters: *α* = 8, *c*_0_ = 0.2. **(C)** Patterns formed under different *α* and *c*_0_ values at *τ* = 10 (bottom) and *τ* = 1000 (top) at fixed *β* = 2. **(D)** Quantities, widths and distances of stripes formed under *β* = 2 at different time points. **(E)** *y*-axis profiles of simulation results under different *α* and *c*_0_ values. The plotted dependent variable *µ*_c_ is the mean concentration along the *x*-axis for a specific *y* position.

The speed of stripe condensation is increased when the enhancement factor *β* is higher (Fig. 5B), consistent with the faster patterning at greater isotropic attraction strengths (Supplementary Fig. S2A). Remarkably, biases as low as *β* = 1.2 suffice to transform isotropic spot patterns into stripes.

We then set the bias to *β* = 2, placing the system firmly in the stripy regime, and investigated the impact of *α* and *c*_0_ on the pattern morphology (Fig. 5C,D). Over time, the number of stripes decreases due to coarsening, analogous to the behavior of the isotropic model (Fig. 2E). The stripes grow in width, accompanied by the narrowing of distances between them as the initial concentration *c*_0_ is increased. However, an effect of the attraction strength *α* on the pattern morphology is not apparent.

In Fig. 5E we show the stripe maturation process. We observed an increase in intensity of individual concentration peaks at earlier stages from *τ* = 10 to *τ* = 100 and a merger into wider condensates throughout *τ* ∈ [10, 1000]. The attraction strength *α* had a small impact on the intensity profiles of stripe patterns early on (*τ* = 10), where a higher *α* is more likely to define sharper stripe boundaries, but its effects are attenuated as time passes by.

#### Anisotropic expansion of an attraction zone

As noted by Murray [52], a narrow tissue domain can force patterns to become essentially one-dimensional. This inspired our second way of implementing anisotropy in the model, a narrow attraction zone with initial dimensions *x*_lim_ and *y*_lim_ that widens over time at rate *v*_exp_ (Fig. 6A). The attraction strength outside of the attraction zone is set to zero (*α* _out_ = 0) and the diffusion rate outside of it is assumed much smaller than within it (*d*_out_ ≪ *d*_in_) to effectively restrict cellular motion to the zone.

**Figure 6:**
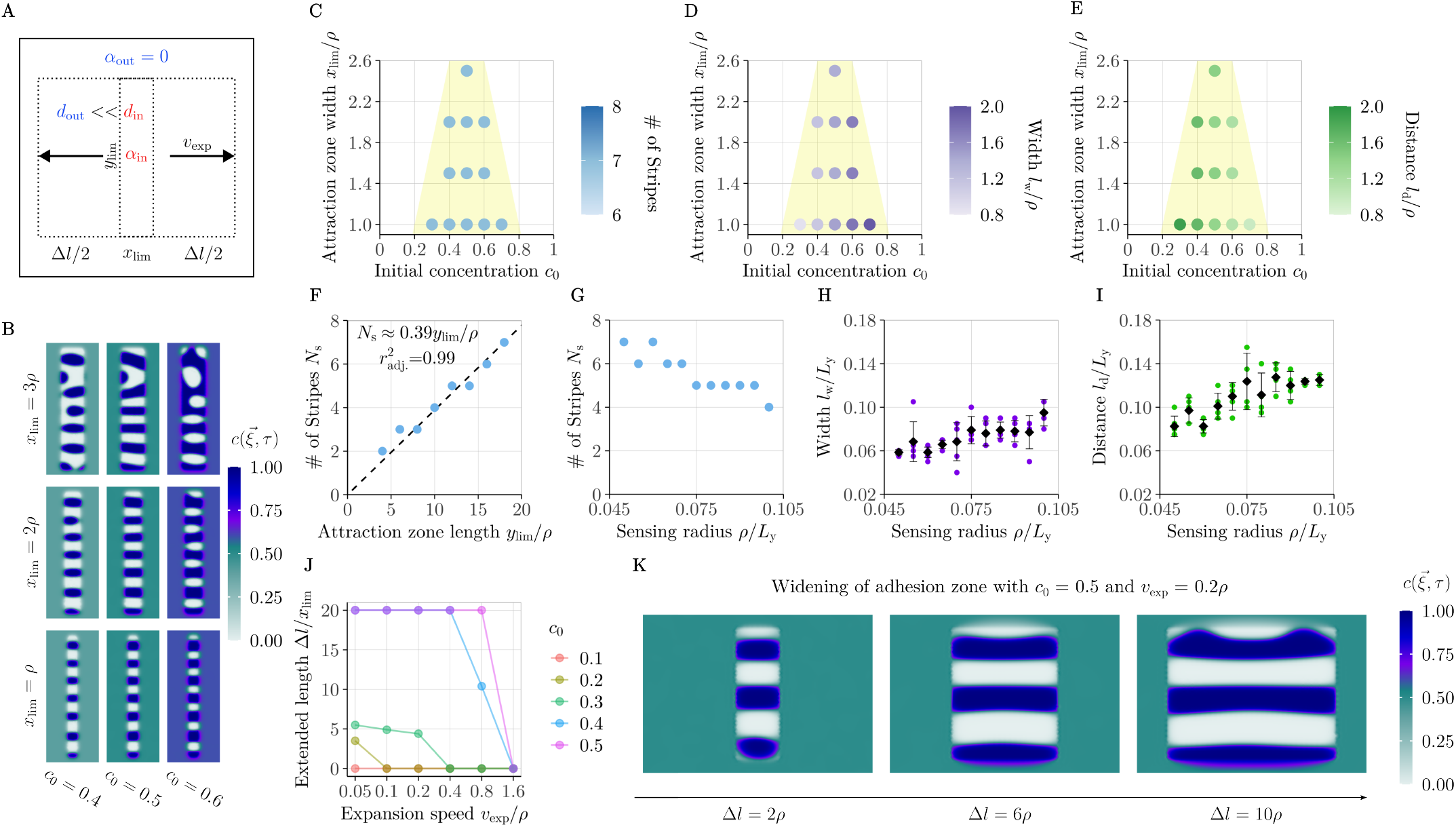
Stripe formation in a widening attraction zone. **(A)** Schematic illustration of the simulation settings. The width of the attraction zone is initially *x*_lim_, and widens at a speed of *v*_exp_. The zone height is *y*_lim_. The diffusivity modules in and out of the attraction zone were set to *d*_in_ = 1 + *d*_out_, *d*_out_ = 0.01. DCM is restricted to the inside of the attraction zone (*α*_out_ = 0). **(B)** Patterns formed under different *c*_0_ and *x*_lim_ values at *τ* = 10. Simulation parameters: *y*_lim_ = 18*ρ, α* = 14. **(C–E)** Quantities, widths and distances of stripes formed under different *c*_0_ and *x*_lim_. The yellow-shaded region schematically indicates the approximate region of stripe formation. Simulation parameters: *y*_lim_ = 18*ρ, α* = 14. **(F)** Impact of the domain height on the number of stripes formed. Blue dots represent numerical results; dashed line is a linear regression. Simulation parameters: *x*_lim_ = 2*ρ, α* = 14, *c*_0_ = 0.4. **(G–I)** Quantities, widths and distances of stripes formed under different sensing radii. *L*_y_ is the *y*-directional length of the computational domain. For widths and distances, colored dots indicate individual measurements; black diamonds and whiskers show the mean and standard deviation. Simulation parameters: *y*_lim_ = 0.9*L*_y_, *L*_y_ = 20, *α* = 14, *c*_0_ = 0.4. (**J**) The effect of initial concentration and expansion rate on the relative increased width Δ*l/x*_lim_ of the attraction zone until the longest stripe no longer spans 80% of the width of the box *x*_lim_ + Δ*l*. The initial zone width is *x*_lim_ = 0.5*ρ*. **(K)** Evolution of stripe expansion under *v*_exp_ = 0.2*ρ, c*_0_ = 0.5 and *x*_lim_ = 0.5*ρ*.

We started by investigating properties of a static attraction zone (*v*_exp_ = 0). Simulation results suggest that stripy patterns can form when the attraction zone is narrow, such as once or twice the sensing radius (Fig. 6B). Furthermore, the further away from 0.5 the initial concentration *c*_0_ is, the less likely domain-spanning stripes are to form (Fig. 6B–E). Once stripes have formed, the width of the attraction zone *x*_lim_ does not have any strong further effect on the quantity, width or separation between stripes given a fixed *y*_lim_. The initial concentration does not affect the number of stripes either, but a higher *c*_0_ increases their width while narrowing the gap between them (Fig. 6C– E). As expected for neighborhood sensing with fixed range, the number of stripes grows linearly with the length of the attraction zone, *y*_lim_ (Fig. 6F). Conversely, tuning the sensing radius is a way of altering the stripe characteristics: Increasing it results in a reduction in stripe number (Fig. 6G), while simultaneously expanding their width and separation (Fig. 6H–I).

Next we turned to a widening attraction zone (*v*_exp_ *>* 0) starting from a initial width of *x*_lim_ = 0.5*ρ*, testing a series of expansion speeds *v*_exp_ and initial concentrations *c*_0_. We recorded the relative change in the expanded zone width Δ*l/x*_lim_ (know as Cauchy strain in mechanics) for which the longest stripe no longer spans across 80% of the attraction zone (Fig. 6J). Δ*l/x*_lim_ = 0 indicates that no such stripes formed at all, and Δ*l/x*_lim_ = 20 represents fully extend stripes spanning the computational domain of width *L*_x_ = 20. Higher expansion speeds resulted in shorter stripes, likely because aggregation cannot keep up with the growth in tissue area that engages in patterning, which leads to patchy stripe formation. The closer the initial concentration *c*_0_ was to 0.5, the longer stripy patterns were preserved, and the more robust they were to increasing expansion rates.

Fig. 6K shows representative stills from the widening process. Recall that in the isotropic version of Model 2, *c*_0_ = 0.5 gives rise to labyrinthine patterns (Fig. 2B), suggesting that this balance between active and inactive cells generally favors stripe formation. These results also demonstrate that a “pre-patterning” process, here induced by the narrow initial attraction zone, can orient patterns in an otherwise fully isotropic model.

## Discussion

Motion-based cellular patterning processes play an important role in biological development [11]. In this article, we reinterpreted and generalized previous differential cell adhesion (DCA) models into a unified directed cell migration (DCM) model covering a much broader range of cellular interactions beyond adhesive forces. The core ingredient is a non-local (integral) term that directs active cellular motion along spatial variations of cellular density, agnostic of the specific biological implementation, be it through differential adhesion, chemotaxis, haptotaxis etc.

Discrete DCA models have given valuable insight on cellular dynamics in the past, but can be computationally demanding and are challenging to study analytically [34, 53, 54]. Continuum models, on the other hand, have allowed to study patterning processes on larger scales and enabled analytical treatment [36–38, 40, 55]. We have extended these studies by quantitatively determining the impact of model parameters on pattern morphology, speed, size and conditions for emergence. While conditions and speed of patterning are determined jointly by all parameters involved, the morphology depends mostly on the initial concentration of active cells, aggregate boundary sharpness mainly on the attraction strength, and the number and size of aggregates on the cells’ sensing radius. By incorporating growing domains and demonstrating mechanisms to orient patterns, we have integrated the DCM model into a biologically relevant context. Furthermore, by implementing the model in a finite element framework, we extended its applicability to 3D, enabling comparisons with real biological morphogenesis.

The coefficient 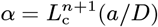 can be understood as Péclet’s number, describing the advective-to-diffusive transport rate ratio. Patterning only occurs with a sufficiently large *α*, indicating that aggregates form when attraction dominates over diffusion. To estimate physiologically plausible *α* values, measurements of the diffusion rate *D* and attraction strength *a* are required. While *D* is measurable experimentally [56], determining *a* is more complex. Although previous studies have preserved accurate ratios of self- and cross-adhesiveness in multi-population DCA models based on experiments [37, 38], direct measurement of *a* with units [m^1*−n*^s^*−*1^] remains unlikely. However, *a* can be indirectly estimated through measurements of cellular speed, 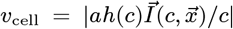, using classical parameter estimation techniques like maximum likelihood [57].

Our DCM model allows for a wide range of interpretations for the advection term in Eq. 1. Unlike previous continuum DCA models, which attribute cellular motion to differential cell adhesion [36–38], our unified DCM framework is largely agnostic of the type of force that gives rise to the directed motion, of which there are many [58]. It also accommodates other mechanisms such as chemotaxis (guided by diffusing molecules), haptotaxis (surface-bound molecules), durotaxis (mechanical properties), topotaxis (topographical cues), and galvanotaxis (electric fields) [59]. Moreover, although originally expressed in this sense, the migrating agents (whose local density is represented by 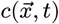 do not necessarily need to be individual cells, but could also be interpreted as cell clumps or aggregates to model collective cell migration [60, 61].

Different forms of migration could also be related to local cell density. For example, recent studies revealed chemotaxis toward the autoinducer AI-2 in E. coli mediated by quorum sensing and diffusing chemicals, which would affect the parameter *a* in our model, influencing patterning time across orders of magnitude (Fig. 3A) [62, 63]. Compared to Turing mechanisms, where patterning speed is tightly linked to molecular diffusion and reaction rates [12] and primarily set by the smaller of the two diffusivities, the DCM model allows for faster patterning even when the cellular diffusivity is lower than that of a morphogen: The distance traveled through directed motility grows linearly with time and thus quickly exceeds that covered by random motion, which grows only with the square root. Future studies could explore additional complexities, such as positional information from morphogen gradients, and more nuanced forms of *w*(*r*) (possibly including repulsion, *w*(*r*) *<* 0) and of the modulators *d*(*c*), *g*(*c*), *h*(*c*). We focused here on simple expressions, but note that other forms are possible. For instance, using *h*(*c*) = (*c*− *c*_1_)(*c*_2_ − *c*)*/*(*c*_2_*− c*_1_)^2^ restricts the range of concentration within which the pattern forms to *c* ∈ [*c*_1_, *c*_2_], rather than spanning the full range [0, 1] as showcased here.

How cell density sensing within a neighborhood of radius *R* is biologically realized may vary, but is generally an open problem. A plausible candidate mechanism is via cytonemes or filopodia. In *Drosophila* wing disc cells, cytonemes can be remarkably straight and reach up to 700 µm in length [64], whereas lengths up to about 40 µm were measured in the air sac primordium [65]. Depending on the tissue and organism, also filopodia can reach dozens or even hundreds of microns in length [66].

A second question that requires further clarification is how cells may translate local density patterns into cell fate specification. That differentiation can be density-dependent has been observed in human mesenchymal stem cells [67]. It has been suggested that differences in cell density may confer differences in cell shape, which is then used as a commitment cue [68].

An interesting parallel may be drawn between the present DCM model and that of active fluids or gels that were recently used to model follicle patterning [23, 24, 69]. In a low-Reynolds regime, viscous stresses within the cells and the surrounding ECM are thought to give rise to a characteristic length scale 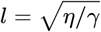 (with a viscosity parameter *η* and a friction coefficient *γ* for sliding against the environment) over which the velocity field decays from a local active stress gradient [70]. Mathematically, this leads to a diffusion-advection equation with a advection velocity *u* whose active component depends on the local density gradient *c*_*x*_: *u* = *l*^2^*u*_*xx*_ + *Ac*_*x*_ [24]. The general form of the DCM (Eq. 3) is similar in its effect: Environmental information is integrated over a spatial region of characteristic extent (here radius *R*), resulting in directional migration. The attraction strength *α* modulates cell activity in a similar way as the coupling coefficient *A* in the active fluid models. In some sense, the DCM model, in its integro-differential form, is more general and might provide a means to biologically interpret the transduction of a non-uniform environment into directed motion without mathematically requiring local gradient information.

Our implementation could be adapted to system-specific characteristics in multiple ways. We constantly observed pattern coarsening in our simulations due to Ostwald ripening or coalescence. Pattern coarsening is also observed in active fluid models [24]. The tendency of aggregates to coalesce into a single structure in the steady state hinders the formation of complex patterns during development. This limitation could be mitigated by incorporating inhibitory effects into the model to stabilize patterns after a defined time period.

Another potential mechanism for spatial separation could involve jamming effects, where cellular structures transition from a fluid-like to a rigid state. This stiffening effect, explained by increased cell density [71], limits cell motility and could be captured within our *d*(*c*) module. Fluid-to-solid transitions may also arise from density-independent factors such as cortical tension and cell-cell adhesion strength [72]. Furthermore, differential growth between two tissue layers could induce the formation of buckled structures [18]. These periodic buckles could act as wells to trap aggregates. Incorporating these neglected effects into our current model could provide a more accurate explanation for some well-spaced patterns observed in biological systems.

We proposed two mechanisms to orient patterns in a certain direction. One method is assuming a narrow but widening zone for diffusion and attraction, the other is directly introducing anisotropy in model parameters. Shoji *et al*. have illustrated the possibility of anisotropic diffusion leading to the orientation of stripes under the biological context that the orientation of scales on fish might cause the directional difference in diffusion [50, 73]. In this study, we implemented anisotropy in the advection term. Either reducing random motility or increasing attraction strength in one direction results in patterns orientated in that direction. A 20% directional bias suffices to orient the stripes. Adhesive forces, for example, could be exerted in an anisotropic manner, as *α*-catenin, which is known to mediate cell-cell adhesion via adherens junction formation, could be activated in cells in an anisotropic manner to exert directed forces [74]. Anisotropy could also arise from anisotropic growth by generating a directed flow field [51].

In summary, the directed cell migration model is a versatile framework for understanding motion-based supracellular patterning during development. With more experimental quantifications of cellular motion likely forthcoming, we anticipate it may explain a variety of developmental patterning systems in future research. In principle, it could also be applied to explain patterning phenomena in different contexts, such as animal swarming, as discussed in [55, 75].

## Materials and Methods

### Numerical simulations

The general integral form of the migratory direction 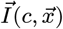 makes the DCM model and its DCA ancestors extraordinarily difficult to solve numerically. Integral terms are not routinely supported by solvers for partial differential equations. All numerical simulations were carried out using the finite element method in COMSOL Multiphysics version 6.2, which provides special operators to carry out the integration discretely. COMSOL model files are provided in the Supplementary Information; a detailed technical description will be provided elsewhere.

The initial condition of the system consisted of a uniformly distributed concentration *c*_0_ with random perturbations (noise) of amplitude *σ*:

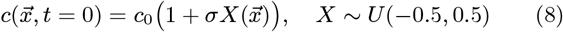

### Linear stability analysis

Critical conditions for pattern formation can be mathematically defined through a linear stability analysis. We start from the non-dimensionalized Eqs. 3 and 4, and approximate 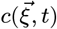 by

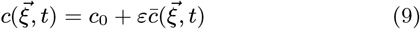

where *c*_0_ is the uniform initial steady-state concentration over an infinite domain that does not vary with space or time, and 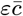 is a small perturbation (*ε* ≪ 1). We can then rewrite the PDE as

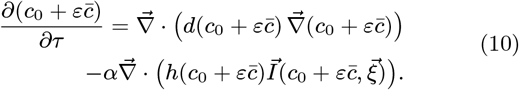

Next we Taylor-expand Eq. 10 to first order about *c*_0_. The Taylor expansion of 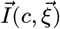 is

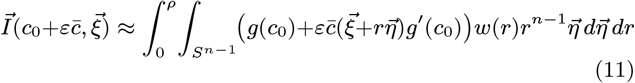

Since 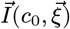 is a symmetric *n*D-spherical integral of a uniform *g*(*c*_0_) multiplied by the directional vector 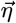it vanishes:

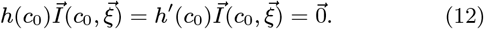

Noting that also 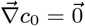 and *∂* _*τ*_ *c*_0_ = 0, Eq. 10 simplifies to

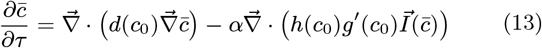

With the usual harmonic ansatz [36, 48]

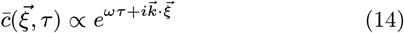

where *ω* is the wave frequency and 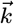 the wave vector, we can then write

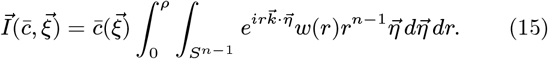

Bringing Eq. 15 back into Eq. 13, we have:

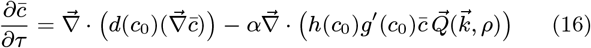

where

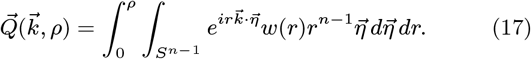

From Eq. 14, it follows that

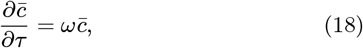

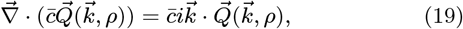

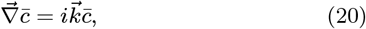

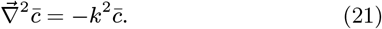

This leads to the dispersion relation

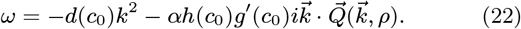

Pattern formation requires the perturbation amplitude to grow, i.e.,

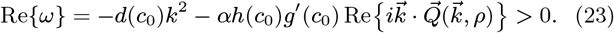

For simplicity, we assume equal weighting of the entire cellular neighborhood (*w* (*r*) ≡ 1) going forward. As (*c*_0_), α, *h* (*c*_0_) and 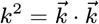 are all positive, 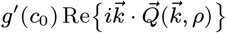 must be negative for Eq. 23 to hold. We can thus rewrite Eq. 23 into

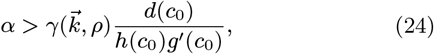

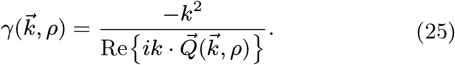

Evaluation of 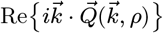 is possible via hyperspherical Bessel functions with hyperspherical Hankel transforms [48], which yields

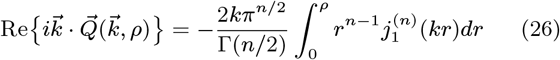

where Γ is the Gamma function and 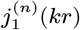 are *n*-dimensional hyperspherical Bessel functions of the first order: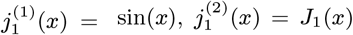, and 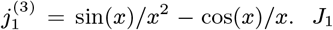 is the Bessel function of the first kind of order one. For 1D, straightforward calculation gives

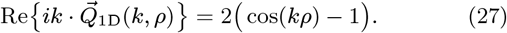

For 2D,

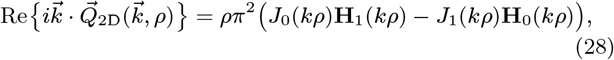

where *J*_0_ is the Bessel function of the first kind of order zero, and **H**_0_ and **H**_1_ are Struve H functions of order zero and one, respectively. For 3D,

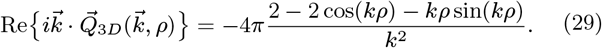

Assuming *g*′(*c*_0_) *>* 0, then 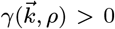 must hold. Critical conditions of *α, c*_0_ and *ρ* appear if we take the smallest positive value of 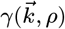, which is assumed as *k →* 0.

For 1D:

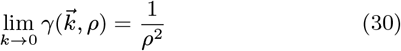

For 2D:

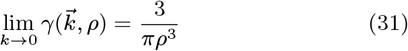

For 3D:

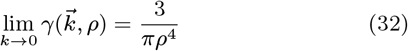

If *g*′(*c*_0_) is negative, such as when *g*(*c*) = *c*(1 − *c*) and *c*_0_ *>* 0.5, this indicates that 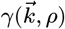 must be negative for patterns to form. In 1D, this is impossible, as *γ* is always positive. In 2D and 3D, *γ* could take negative values. We thus look for the smallest positive value of 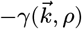, or the largest negative value of 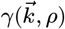. For *ρ* = 1, numerical evaluation yields

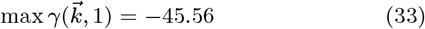

in 2D and

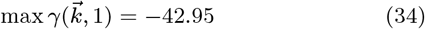

in 3D. Note that this is an over 40-fold upsurge in the critical attraction strength needed for patterns to form compared to *c*_0_ *<* 0.5, which is numerically inaccessible in our implementation, and may also biologically be difficult to achieve.

### Domain growth

To avoid computationally expensive re-meshing in simulations with domain growth, we transformed the coordinate system from the Eulerian to a Lagrangian frame, which enables growth simulations on a static domain. We start from the general form of Eqs. 3 and 4 with a velocity field 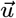 generated by domain growth:

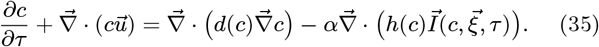

Eq. 35 is transformed into Lagrangian coordinates by setting 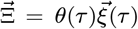 where the stretching factor is *θ*(*τ*) = *L*_0_*/L*(*τ*) and the side length of the uniformly growing domain is *L*(*τ*) = *L*_0_ + *vτ*. We obtain *∂* Ξ _*i*_ */ ∂ ξ* _*i*_ = *θ* (*τ*) in each direction *i* = 1, …, *n*. Under this transformation, the velocity field becomes 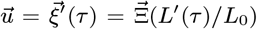. The term 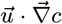 disappears in the Lagrangian framework. The gradient operator transforms as 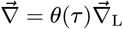. Rewriting 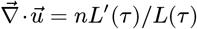 in *n*D and *L*′(*τ*) = *v*, Eq. 35 becomes

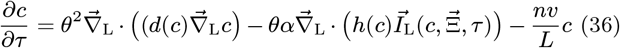

with

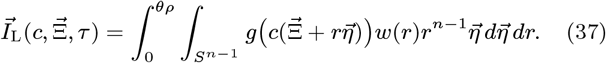

Note that we restrict our analysis to isotropic growth here. The sensing integral 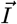 would become an ellipse integral in 2D, or an ellipsoid integral in 3D if the growth was anisotropic.

### Extraction of aggregate features

For all isotropic models, simulation results were first transformed into intensity images and then analyzed using scikit-image [76] in Python. The images were first segregated into fore- and background using a 1.1-fold higher threshold than the threshold calculated in Ostu’s method [77] to separate higher concentrated regions that are bridged by low concentration gaps. Segments with area smaller than 15 pixels were filtered out, as they were mostly noise.

Images produced under periodic boundary conditions needed special processing, as these images contain truncated segments at borders, whose other parts appear on the opposite side. Without processing, these segments would be counted as separate objects. These images were first duplicated and then stitched together in a 3 *×* 3 array. After labeling of the segments, all segments without intersection with the center image tile were filtered out. Next, all segments that touched the borders of the large stitched image were filtered out to exclude domain-spanning (infinite) segments. Finally, a detection of duplicates was performed by only preserving one segment among those that have both the same area and mean intensity. By executing the above steps, we transformed truncated and separate segments into complete objects without repetition. However, note that this does not count any segment that extends across all four borders of the computational domain.

For models with anisotropy that produced stripes, stripe widths and distances were quantified as follows: Intensity images were averaged in the image direction aligned with the stripes, producing mean concentration profiles similar to those shown in Fig. 5E. The maximum *I*_max_ and minimum *I*_min_ of that intensity profile were extracted, and a reference line was set at (*I*_max_ + *I*_min_)*/*2. The stripe width was defined as the distance of intersection points between the reference line and the profile around each peak, whereas the distance between stripes was defined by the distance of intersection points around each trough.

## Acknowledgment

The authors thank Vincent Zanetta and Barbara Walkowiak for preliminary model assessment, Walter Frei and Sven Friedel from COMSOL Inc. for technical assistance with the numerical implementation, as well as the CoBi group at ETH Zürich, Kevin J. Painter and José A. Carrillo for discussions.

## SUPPLEMENTARY FIGURES

**Fig. S1:**
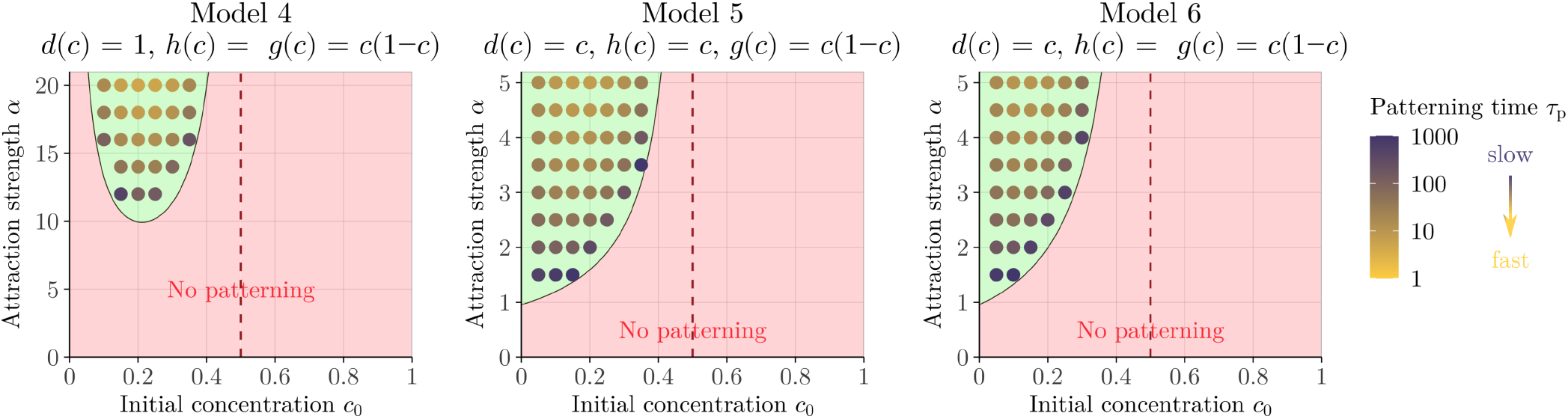
Patterning conditions and speed for further model variants. Phase space of patterning for three further model variants as labeled. Green-shaded areas indicate *α >* 3*d*(*c*_0_)*/*(*h*(*c*_0_)*g*′(*c*_0_)*πρ*^3^). Points represent numerical simulations that produced patterns; absence of a point in the grid indicates no pattern. Colors of points represent the time point of pattern formation, *τ*_p_, as defined in Fig. 3B, from slow (dark purple) to fast (yellow). Simulation settings: *ρ* = 1, *σ* = 0.01.

**Fig. S2:**
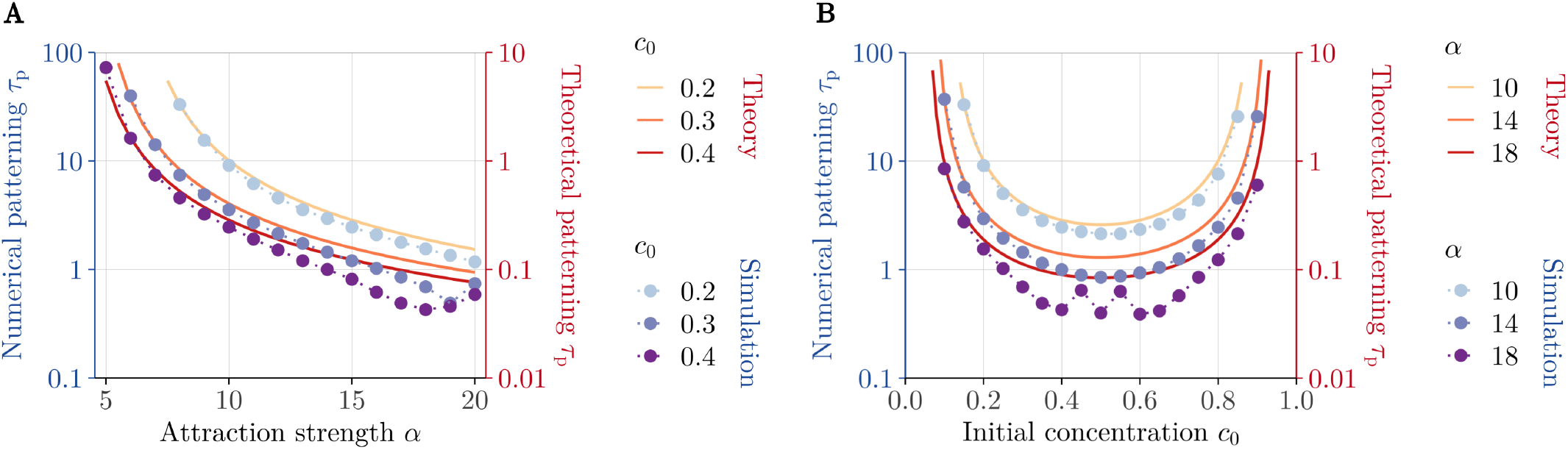
Dependency of the patterning time on the attraction strength and initial concentration. **(A)** Influence of the attraction strength *α* on the numerical (blue color scale) and theoretical (red color scale) patterning time *τ*_p_ for different *c*_0_ values. **(B)** Influence of the initial concentration *c*_0_ on the numerical (blue color scale) and theoretical (red color scale) patterning time *τ*_p_ for different *α* values.

## Notes

### Competing Interest Statement

The authors have declared no competing interest.

## References

[1] A. M. Turing. The chemical basis of morphogenesis. Philos. Trans. R. Soc. Lond. B, 237:37–72, 1952. doi: 10.1098/rstb.1952.0012.

[2] L. Wolpert. Positional information and the spatial pattern of cellular differentiation. J. Theor. Biol., 25:1–47, 1969. doi: 10.1016/S0022-5193(69)80016-0.

[3] J. D. Murray. Mathematical Biology II. Spatial Models and Biomedical Applications. Springer, 2003. doi: 10.1007/b98869.

[4] D. Drasdo. Buckling Instabilities of One-Layered Growing Tissues. Phys. Rev. Lett., 84:4244–4247, 2000. doi: 10.1103/PhysRevLett.84.4244.

[5] A. E. Shyer, T. Tallinen, N. L. Nerurkar, Z. Wei, E. S. Gil, D. L. Kaplan, C. J. Tabin, and L. Mahadevan. Villification: How the Gut Gets Its Villi. Science, 342:212–218, 2013. doi: 10.1126/science.1238842.

[6] T. Hirashima. Pattern Formation of an Epithelial Tubule by Mechanical Instability during Epididymal Development. Cell Rep., 9:866–873, 2014. doi: 10.1016/j.celrep.2014.09.041.

[7] V. D. Varner, J. P. Gleghorn, E. Miller, D. C. Radisky, and C. M. Nelson. Mechanically patterning the embryonic airway epithelium. Proc. Natl. Acad. Sci. U.S.A., 112:9230– 9235, 2015. doi: 10.1073/pnas.1504102112.

[8] F. L. Lampart, R. Vetter, K. A. Yamauchi, Y. Wang, S. Runser, N. Strohmeyer, F. Meer, M.-D. Hussherr, G. Camenisch, H.-H. Seifert, C. A. Rentsch, C. Le Magnen, L. Müller, D. Bubendorf, and D. Iber. Morphometry and mechanical instability at the onset of epithelial bladder cancer. Nat. Phys., 2025. doi: 10.1038/s41567-024-02735-2.

[9] J. Holtfreter. Gewebeaffinität, ein Mittel der embryonalen Formbildung. Arch. Exp. Zellforsch. Gewebeszücht, 23:169– 209, 1939.

[10] Malcolm S Steinberg. Reconstruction of tissues by dissociated cells: some morphogenetic tissue movements and the sorting out of embryonic cells may have a common explanation. Science, 141(3579):401–408, 1963.

[11] T. Fulton, B. Verd, and B. Steventon. The unappreciated generative role of cell movements in pattern formation. R. Soc. Open Sci., 9:211293, 2022. doi: 10.1098/rsos.211293.

[12] Y. Mori, A. Jilkine, and L. Edelstein-Keshet. Wave-pinning and cell polarity from a bistable reaction-diffusion system. Biophys. J., 94:3684–3697, 2008. doi: 10.1529/biophysj.107.120824.

[13] L. Marcon and J. Sharpe. Turing patterns in development: what about the horse part? Curr. Opin. Genet. Dev., 22: 578–584, 2012. doi: 10.1016/j.gde.2012.11.013.

[14] M. Mercker, F. Brinkmann, A. Marciniak-Czochra, and T. Richter. Beyond turing: mechanochemical pattern formation in biological tissues. Biol. Direct., 11:2, 2016. doi: 10.1186/s13062-016-0124-7.

[15] D. Iber, M. Mederacke, and R. Vetter. Coordination of nephrogenesis with branching of the urinary collecting system, the vasculature and the nervous system, volume 163, chapter 3, pages 45–82. Academic Press, 2025. doi: 10.1016/bs.ctdb.2024.11.008.

[16] T. Kurics, D. Menshykau, and D. Iber. Feedback, receptor clustering, and receptor restriction to single cells yield large turing spaces for ligand-receptor-based turing models. Phys. Rev. E, 90:022716, 2014. doi: 10.1103/PhysRevE.90.022716.

[17] D. Menshykau, P. Blanc, E. Unal, V. Sapin, and D. Iber. An interplay of geometry and signaling enables robust lung branching morphogenesis. Development, 141:4526–4536, 2014. doi: 10.1242/dev.116202.

[18] Biot M. A. Folding instability of a layered viscoelastic medium under compression. Proc. R. Soc. Lond. A, 242: 444–454, 1957. doi: 10.1098/rspa.1957.0187.

[19] D. P. Richman, R. M. Stewart, J. Hutchinson, and V. S. Caviness. Mechanical Model of Brain Convolutional Development. Science, 189:18–21, 1975. doi: 10.1126/science.1135626.

[20] R. A. Foty and M. S. Steinberg. Differential adhesion in model systems. WIREs Dev. Biol., 2:631–645, 2013. doi: 10.1002/wdev.104.

[21] T. Y.-C. Tsai, R. M. Garner, and S. G. Megason. Adhesion-Based Self-Organization in Tissue Patterning. Annu. Rev. Cell Dev. Biol., 38:349–374, 2022. doi: 10.1146/annurev-cellbio-120420-100215.

[22] D. Iber and M. Mederacke. Tracheal Ring Formation. Front. Cell Dev. Biol., 10:900447, 2022. doi: 10.3389/fcell.2022.900447.

[23] A. E. Shyer, A. R. Rodrigues, G. G. Schroeder, E. Kassianidou, S. Kumar, and R. M. Harland. Emergent cellular self-organization and mechanosensation initiate follicle pattern in the avian skin. Science, 357:811–815, 2017. doi: 10.1126/science.aai7868.

[24] K. H. Palmquist, S. F. Tiemann, F. L. Ezzeddine, S. Yang, C. R. Pfeifer, A. Erzberger, A. R. Rodrigues, and A. E. Shyer. Reciprocal cell-ECM dynamics generate supracellular fluidity underlying spontaneous follicle patterning. Cell, 185:1960–1973, 2022. doi: 10.1016/j.cell.2022.04.023.

[25] B. Sorre, A. Warmflash, A. H. Brivanlou, and E. D. Siggia. Encoding of temporal signals by the TGF-beta pathway and implications for embryonic patterning. Dev. Cell, 30: 334–342, 2014. doi: 10.1016/j.devcel.2014.05.022.

[26] H. M. McNamara, S. C. Solley, M. M. Adamson, B. an Chan, and J. E. Toettcher. Recording morphogen signals reveals mechanisms underlying gastruloid symmetry breaking. Nat. Cell Biol., 26:1832–1844, 2024. doi: 10.1038/s41556-024-01521-9.

[27] S. Gsell, S. Tlili, M. Merkel, and P.-F. Lenne. Marangoni-like tissue flows enhance symmetry breaking of embryonic organoids. Nat. Phys., 21:644–653, 2025. doi: 10.1038/s41567-025-02802-2.

[28] J. Adler. Chemoreceptors in Bacteria. Science, 166:1588– 1597, 1969. doi: 10.1126/science.166.3913.1588.

[29] S. B. Carter. Principles of Cell Motility: The Direction of Cell Movement and Cancer Invasion. Nature, 208:1183– 1187, 1965. doi: 10.1038/2081183a0.

[30] S. B. Carter. Haptotaxis and the Mechanism of Cell Motility. Nature, 213:256–260, 1967. doi: 10.1038/213256a0.

[31] C.-M. Lo, H.-B. Wang, M. Dembo, and Y.-I. Wang. Cell Movement Is Guided by the Rigidity of the Substrate. Biophys. J., 79:144–152, 2000. doi: 10.1016/S0006-3495(00)76279-5.

[32] J. Park, D.-H. Kim, and A. Levchenko. Topotaxis: A New Mechanism of Directed Cell Migration in Topographic ECM Gradients. Biophys. J., 114:1257–1263, 2018. doi: 10.1016/j.bpj.2017.11.3813.

[33] M. S. Steinberg. Does differential adhesion govern self-assembly processes in histogenesis? Equilibrium configurations and the emergence of a hierarchy among populations of embryonic cells. J. Exp. Zool., 173:395–433, 1970. doi: 10.1002/jez.1401730406.

[34] J. A. Glazier and F. Graner. Simulation of the differential adhesion driven rearrangement of biological cells. Phys. Rev. E, 47:2128, 1993. doi: 10.1103/physreve.47.2128.

[35] J. D. Murray and G. F. Oster. Cell traction models for generating pattern and form in morphogenesis. J. Math. Biol., 19:265–279, 1984. doi: 10.1007/BF00277099.

[36] N. J. Armstrong, K. J. Painter, and J. A. Sherratt. A continuum approach to modelling cell-cell adhesion. J. Theor. Biol., 243:98–113, 2006. doi: 10.1016/j.jtbi.2006.05.030.

[37] H. Murakawa and H. Togashi. Continuous models for cell–cell adhesion. J. Theor. Biol., 374:1–12, 2015. doi: 10.1016/j.jtbi.2015.03.002.

[38] J. A. Carrillo, H. Murakawa, M. Sato, H. Togashi, and O. Trush. A population dynamics model of cell-cell adhesion incorporating population pressure and density saturation. J. Theor. Biol., 474:14–24, 2019. doi: 10.1016/j.jtbi.2019.04.023.

[39] T. Sekimura, M. Zhu, J. Cook, P. K. Maini, and J. D. Murray. Pattern formation of scale cells in lepidoptera by differential origin-dependent cell adhesion. Bull. Math. Biol., 61:807–828, 1999. doi: 10.1006/bulm.1998.0062.

[40] K. J. Painter, T. Hillen, and J. R. Potts. Biological modeling with nonlocal advection–diffusion equations. Math. Models Methods Appl. Sci., 34:57–107, 2024. doi: 10.1142/S0218202524400025.

[41] A. Gerisch. On the approximation and efficient evaluation of integral terms in PDE models of cell adhesion. IMA J. Numer. Anal., 30:173–194, 10 2009. doi: 10.1093/imanum/drp027.

[42] A. Gerisch and K. J. Painter. Mathematical Modeling of Cell Adhesion and Its Applications to Developmental Biology and Cancer Invasion. In A. Chauviere, L. Preziosi, and C. Verdier, editors, Cell Mechanics: From Single Scale-Based Models to Multiscale Modeling, chapter 12. Chapman and Hall/CRC, 2010. doi: 10.1201/9781420094558.

[43] I. Aziz and I. Khan. Numerical Solution of Diffusion and Reaction–Diffusion Partial Integro-Differential Equations. Int. J. Comput. Meth., 15:1850047, 2018. doi: 10.1142/S0219876218500470.

[44] A. Gerisch and M.A.J. Chaplain. Mathematical modelling of cancer cell invasion of tissue: Local and non-local models and the effect of adhesion. J. Theor. Biol., 250:684–704, 2008. doi: 10.1016/j.jtbi.2007.10.026.

[45] C. Cui. Dynamics of cell movement and tissue motion in gastrulation and micromass cell culture. PhD thesis, Indiana University, 2005. URL https://infomall.org/sites/dsc/biocomplexity/jglazier/docs/student_dissertations/cuithesis.pdf.

[46] J. S. Bois, F. Jülicher, and S. W. Grill. Pattern Formation in Active Fluids. Phys. Rev. Lett., 106:028103, 2011. doi: 10.1103/PhysRevLett.106.028103.

[47] P. Taylor. Ostwald ripening in emulsions. Adv. Colloid Interface Sci., 75:107–163, 1998. doi: 10.1016/S0001-8686(98)00035-9.

[48] T. J. Jewell, A. L. Krause, P. K. Maini, and E. A. Gaffney. Patterning of nonlocal transport models in biology: The impact of spatial dimension. Math. Biosci., 366:109093, 2023. doi: 10.1016/j.mbs.2023.109093.

[49] S. Yin and L. Mahadevan. Contractility-Induced Phase Separation in Active Solids. Phys. Rev. Lett., 131:148401, 2023. doi: 10.1103/PhysRevLett.131.148401.

[50] H. Shoji, Y. Iwasa, A. Mochizuki, and S. Kondo. Directionality of Stripes Formed by Anisotropic Reaction–Diffusion Models. J. Theor. Biol., 214:549–561, 2002. doi: 10.1006/jtbi.2001.2480.

[51] T. W. Hiscock and S. G. Megason. Orientation of Turing-like Patterns by Morphogen Gradients and Tissue Anisotropies. Cell Syst., 1:408–416, 2015. doi: 10.1016/j.cels.2015.12.001.

[52] J. D. Murray. How the Leopard Gets Its Spots. Sci. Am., 258:80–87, 1988.

[53] D. Sulsky, S. Childress, and J. K. Percus. A model of cell sorting. J. Theor. Biol., 106:275–301, 1984. doi: 10.1016/0022-5193(84)90031-6.

[54] F. Graner and J. A. Glazier. Simulation of biological cell sorting using a two-dimensional extended potts model. Phys. Rev. Lett., 69:2013, 1992. doi: 10.1103/PhysRevLett.69.2013.

[55] K. J. Painter, J. M. Bloomfield, J. A. Sherratt, and A. Gerisch. A nonlocal model for contact attraction and repulsion in heterogeneous cell populations. Bull. Math. Biol., 77:1132–1165, 2015. doi: 10.1007/s11538-015-0080-x.

[56] A. Kitamura and M. Kinjo. Determination of diffusion coefficients in live cells using fluorescence recovery after photobleaching with wide-field fluorescence microscopy. Biophysics and Physicobiology, 15:1–7, 2018. doi: 10.2142/biophysico.15.01.

[57] T. Glimm, B. KaŚmierczak, C. Cui, S. A. Newman, and R. Bhat. Precartilage condensation during limb skeletogenesis occurs by tissue phase separation controlled by a bistable cell-state switch with suppressed oscillatory dynamics. bioRxiv, 2021. doi: 10.1101/2021.03.15.435301.

[58] I. C. Fortunato and R. Sunyer. The Forces behind Directed Cell Migration. Biophysica, 2:548–563, 2022. doi: 10.3390/biophysica2040046.

[59] S. SenGupta, C. A. Parent, and J. E. Bear. The principles of directed cell migration. Nat. Rev. Mol. Cell Biol., 22: 529–547, 2021. doi: 10.1038/s41580-021-00366-6.

[60] P. Friedl and D. Gilmour. Collective cell migration in morphogenesis, regeneration and cancer. Nat. Rev. Mol. Cell Biol., 10:445–457, 2009. doi: 10.1038/nrm2720.

[61] A. Haeger, K. Wolf, M. M. Zegers, and P. Friedl. Collective cell migration: guidance principles and hierarchies. Trends Cell Biol., 25:556–566, 2015. doi: 10.1016/j.tcb.2015.06.003.

[62] L. Laganenka, J. Lee, L. Malfertheiner, C. L. Dieterich, L. Fuchs, J. Piel, C. von Mering, V. Sourjik, and W. Hardt. Chemotaxis and autoinducer-2 signalling mediate colonization and contribute to co-existence of Escherichia coli strains in the murine gut. Nat. Microbiol., 8:204–217, 2023. doi: 10.1038/s41564-022-01286-7.

[63] M. B. Miller and B. L. Bassler. Quorum Sensing in Bacteria. Ann. Rev. Microbiol., 55:165–199, 2001. doi: 10.1146/annurev.micro.55.1.165.

[64] F.-A. Ramírez-Weber and T. B. Kornberg. Cytonemes: Cellular Processes that Project to the Principal Signaling Center in Drosophila Imaginal Discs. Cell, 97:599–607, 1999. doi: 10.1016/S0092-8674(00)80771-0.

[65] S. Roy, H. Huang, S. Liu, and T. B. Kornberg. Cytoneme-Mediated Contact-Dependent Transport of the Drosophila Decapentaplegic Signaling Protein. Science, 343:1244624, 2014. doi: 10.1126/science.1244624.

[66] H. T. Hu, T. Nishimura, H. Kawana, R. A. S. Dante, G. D’Angelo, and S. Suetsugu. The cellular protrusions for inter-cellular material transfer: similarities between filopodia, cytonemes, tunneling nanotubes, viruses, and extra-cellular vesicles. Front. Cell Dev. Biol., 12:1422227, 2024. doi: 10.3389/fcell.2024.1422227.

[67] M. F. Pittenger, A. M. Mackay, S. C. Beck, R. K. Jaiswal, R. Douglas, J.D. Mosca, M. A. Moorman, D. W. Simonetti, S. Craig, and D. R. Marshak. Multilineage Potential of Adult Human Mesenchymal Stem Cells. Science, 284:143– 147, 1999. doi: 10.1126/science.284.5411.143.

[68] R. McBeath, D. M. Pirone, C. M. Nelson, K. Bhadriraju, and C. S. Chen. Cell Shape, Cytoskeletal Tension, and RhoA Regulate Stem Cell Lineage Commitment. Dev. Cell, 6:483–495, 2004. doi: 10.1016/S1534-5807(04)00075-9.

[69] S. Yang, K. H. Palmquist, L. Nathan, C. R. Pfeifer, P. J. Schultheiss, A. Sharma, L. C. Kam, P. W. Miller, A. E. Shyer, and A. R. Rodrigues. Morphogens enable interacting supracellular phases that generate organ architecture. Science, 382:eadg5579, 2023. doi: 10.1126/science.adg5579.

[70] M. Mayer, M. Depken, J. Bois, F. Jülicher, and S. W. Grill. Anisotropies in cortical tension reveal the physical basis of polarizing cortical flows. Nature, 467:617–621, 2010. doi: 10.1038/nature09376.

[71] T. E. Angelini, E. Hannezo, X. Trepat, M. Marquez, J. J. Fredberg, and D. A. Weitz. Glass-like dynamics of collective cell migration. Proc. Natl. Acad. Sci. U.S.A., 108: 4714–4719, 2011. doi: 10.1073/pnas.1010059108.

[72] D. Bi, J. H. Lopez, J. M. Schwarz, and M. L. Manning. A density-independent rigidity transition in biological tissues. Nat. Phys., 11:1074–1079, 2015. doi: 10.1038/nphys3471.

[73] H. Shoji, A. Mochizuki, Y. Iwasa, M. Hirata, T. Watanabe, S. Hioki, and S. Kondo. Origin of directionality in the fish stripe pattern. Dev. Dynam., 226:627–633, 2003. doi: 10.1002/dvdy.10277.

[74] K. Matsuzawa, T. Himoto, Y. Mochizuki, and J. Ikenouchi. α-Catenin Controls the Anisotropy of Force Distribution at Cell-Cell Junctions during Collective Cell Migration. Cell Rep., 23:3447–3456, 2018. doi: 10.1016/j.celrep.2018.05.070.

[75] J. A. Carrillo, R. Eftimie, and F. Hoffmann. Non-local kinetic and macroscopic models for self-organised animal aggregations. Kinet. Relat. Models, 8:413–441, 2015. doi: 10.3934/krm.2015.8.413.

[76] S. Van der Walt, J. L. Schönberger, J. Nunez-Iglesias, F. Boulogne, J. D. Warner, N. Yager, E. Gouillart, and T. Yu. scikit-image: image processing in Python. PeerJ, 2: e453, 2014. doi: 10.7717/peerj.453.

[77] N. Otsu. A Threshold Selection Method from Gray-Level Histograms. IEEE Trans. Syst. Man. Cybern., 9:62–66, 1979. doi: 10.1109/TSMC.1979.4310076.

